# Silencing cathepsin L expression reduces *Myzus persicae* protein content and the nutritional value as prey for *Coccinella septempunctata*

**DOI:** 10.1101/451211

**Authors:** Imran Rauf, Muhammad Asif, Imran Amin, Rubab Zahra Naqvi, Noroza Umer, Shahid Mansoor, Georg Jander

**Author notes:** To whom correspondence should be addressed: Georg Jander, Boyce Thompson Institute, 533 Tower Road, Ithaca NY 14853, USA.

## Abstract

Gut-expressed aphid genes, which may be more easily inhibited by RNA interference (RNAi) constructs, are attractive targets for pest control efforts involving transgenic plants. Here we show that expression of cathepsin L, a cysteine protease that functions in aphid guts, can be reduced by expression of an RNAi construct in transgenic tobacco. The effectiveness of this approach is demonstrated by up to 80% adult mortality, reduced fecundity, and delayed nymph production of *Myzus persicae* (green peach aphids) when cathepsin L expression was reduced by plant-mediated RNAi. Consistent with the function of cathepsin L as a gut protease, *M. persicae* fed on the RNAi plants had a lower protein content in their bodies and excreted more protein in their honeydew. Larvae of *Coccinella septempunctata* (seven-spotted ladybugs) grew more slowly on aphids having reduced cathepsin L expression, suggesting that prey insect nutritive value, and not just direct negative effects of the RNAi construct, needs to be considered when producing transgenic plants for RNAi-mediated pest control.

**Highlights:**

- Silencing expression of cathepsin L by RNA interference reduces protein content of *Myzus persicae* (green peach aphid) bodies.
- Honeydew of aphids with cathepsin L silenced contains elevated protein.
- Cathepsin L is required for efficient protein uptake from phloem sap.
- Aphids with cathepsin L expression silenced have increased mortality and fewer offspring.
- *Coccinella septempunctata* (seven-spotted ladybugs) grow more slowly on aphids with expression of cathepsin L silenced.

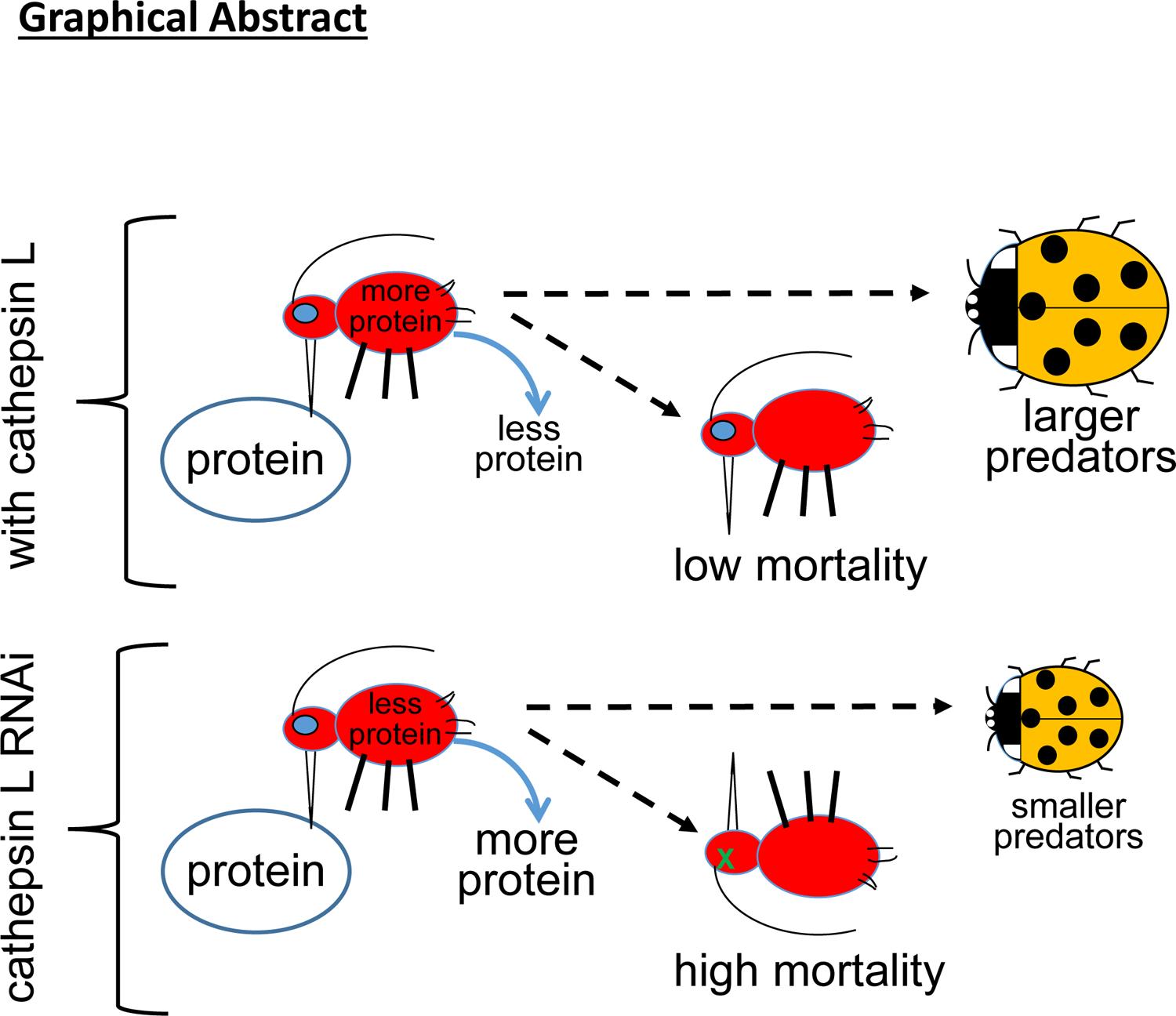

## 1. Introduction

The green peach aphid, *Myzus persicae* (Suzler) (Homoptera: Aphididae), is one of the most deleterious insect pests world-wide. It has a polymorphic appearance and a wide host range, infesting more than 400 species in 40 plant families (Blackman and Eastop, 2000). High dispersive qualities, adaptability to extreme weather conditions, parthenogenetic reproduction, dense populations, and aggregated feeding behavior are some of the key features of this pest that make it an aggressive pest in agricultural systems. Being sap-sucking feeders, *M. persicae* not only inhibit plant growth by removing photoassimilates but also transfer viruses between host plants. Aphids feed by inserting their stylets into the epidermis of the leaves, continue by puncturing mesophyll cells and testing their cytosolic contents, and eventually reach stable feeding sites in the phloem (Powell et al., 2006). Amino acids, in both free and protein-bound form, are among the key nutrients that aphids obtain from phloem tissues (Douglas, 2006).

RNA interference (RNAi), a form of homology-dependent gene expression silencing, is being developed as a management tool for the inhibition of insect pests. In most approaches, RNAi depends on the introduction of target-specific double-stranded RNA (dsRNA) into an organism, which leads to silencing of the target gene by endogenous cleavage pathways (Yu et al., 2016). This technology has been used extensively to reduce gene expression in several insect orders, including Diptera (Li et al., 2011), Lepidoptera (Terenius et al., 2011), Coleoptera (Cao et al., 2018; Wan et al., 2015), Blattodea (Huang and Lee, 2011) and Hemiptera (Bhatia and Bhattacharya, 2018; Cao et al., 2018; Elzinga et al., 2014; Mao and Zeng, 2013; Tzin et al., 2015; Wilson et al., 2011). Methods for introducing dsRNA into target organisms include microinjection, oral feeding from artificial diets with dsRNA (Wang et al., 2015; Yu et al., 2013), and the expression of dsRNA constructs in genetically engineered plants. (Navale P.M. and Krishna V., 2014; Raza et al., 2016; Zha et al., 2011; Zhang et al., 2017). The success of such plant-mediated RNAi often is dependent on the selection of target genes. Based on prior studies, target genes involved in physiology, metamorphosis, metabolic processes, and development can be effective RNAi targets in aphids. Transcriptomic studies have confirmed that genes involved in the digestive process have a high probability of being important for aphid survival (Zhang et al., 2013).

Cysteine proteases, in particular cathepsin L and cathepsin B, are preferentially expressed in aphid guts (Cristofoletti et al., 2003; Deraison et al., 2004; Pyati et al., 2011; Ramsey et al., 2007; Rispe et al., 2008) and are thought to function in the degradation and uptake of phloem proteins, which can be fairly abundant, up to 60 mg/ml in some plants (Cronshaw and Sabnis, 1989; Kehr, 2006). Several lines of evidence indicate an important digestive function for cathepsins, making them attractive targets for plant-mediated RNAi. Barley protease inhibitors fed to *M. persicae* and *Acyrthosiphon pisum* (pea aphid) reduced performance on artificial diet and/or host plants (Carrillo et al., 2011). Similarly, feeding of protease inhibitors from wheat inhibited protein digestion by *Sitobion avenae* (Pyati et al., 2011). In a transgenic approach, expression of oryzacystatin in transgenic *Brassica napus* decreased *M. persicae* biomass by up to 40% (Rahbe et al., 2003). Injection of double-stranded RNA targeting cathepsin L reduced expression of this gut-expressed gene in *Acyrthosiphon pisum* (pea aphid) (Jaubert-Possamai et al., 2007). RNAi-mediated expression silencing of cathepsin B genes inhibited *M. persicae* in host plant-specific manner (Mathers et al., 2017). Feeding *A. pisum* with cathepsin L dsRNA in artificial diet specifically silenced gene expression in the gut, whereas injection of the dsRNA caused more widespread effects (Sapountzis et al., 2014b). Cathepsin activity in the aphid gut also was demonstrated by engineering a cleavage sites for these enzyme into the Bt toxin Cry4Aa to allow activation (Rausch et al., 2016).

Both predators and parasitoids play important roles in the suppression of insect pests. Whereas predatory insects tend to be larger than their prey and need more than one prey item to complete their development, parasitoids are smaller than their hosts, complete their development in a single prey insect, and eventually kill it. It is possible that plant-mediated RNAi would disturb not only the activity and physiology of the targeted insects, but also non-target insects like predators and parasitoids that consume them (Roberts et al., 2015). However, based on the high specificity of the dsRNA and the ability to design species-specific constructs, the chance of adverse effects on predators and parasitoids is presumed to be very low (Bachman et al., 2013).

We have developed a multi-species RNAi based construct designed to knock down expression of cathepsin L from *M. persicae*, ecdysone receptor from *Bemisia tabaci* (whitefly), and chitin synthase from *Phenacoccus solenopsis* (cotton mealybug). The construct was transformed into the model plant *Nicotiana tabacum* (tobacco) for evaluation of gene expression knockdown effects on these three phloem-feeding insect species. Here we describe the specific effects of cathepsin L knockdown on *M. persicae* growth and interactions with the seven-spotted ladybug, *Coccinella septempunctata* (Coleoptera: Coccinellidae), a voracious predator of many aphid species.

## 2. Materials and methods

### 2.1. Development of the RNAi construct

Three genes with essential insect functions, *M. persicae* cathepsin L (AJ496197), *B. tabaci* Ecdysone receptor (EF174329), and *P. solenopsis* chitin synthase (KF384512) were selected for making an RNAi construct. Sequences from different strains/species of the same genus were aligned using multiple alignments, maximum likelihood by Mega 6 software. A highly conserved region (200 bps) of each gene (Table S1) was identified and used for further construct development. An RNAi construct was made with a fusion of the three 200-bp sequences in the sense and antisense orientation. For sense orientation, the 600-bp sequence was synthesized by Eurofins, Genomics, Canada. The sense-orientation sequence was cloned into the pFGC-5941 RNAi vector with XhoI and NcoI restriction sites at 5’ and 3’ respectively. The antisense construct was cloned into pFGC-5941 using the BamHI and XbaI restriction sites at 5’ and 3’ ends, respectively (Fig. 1A).

**Figure 1.**
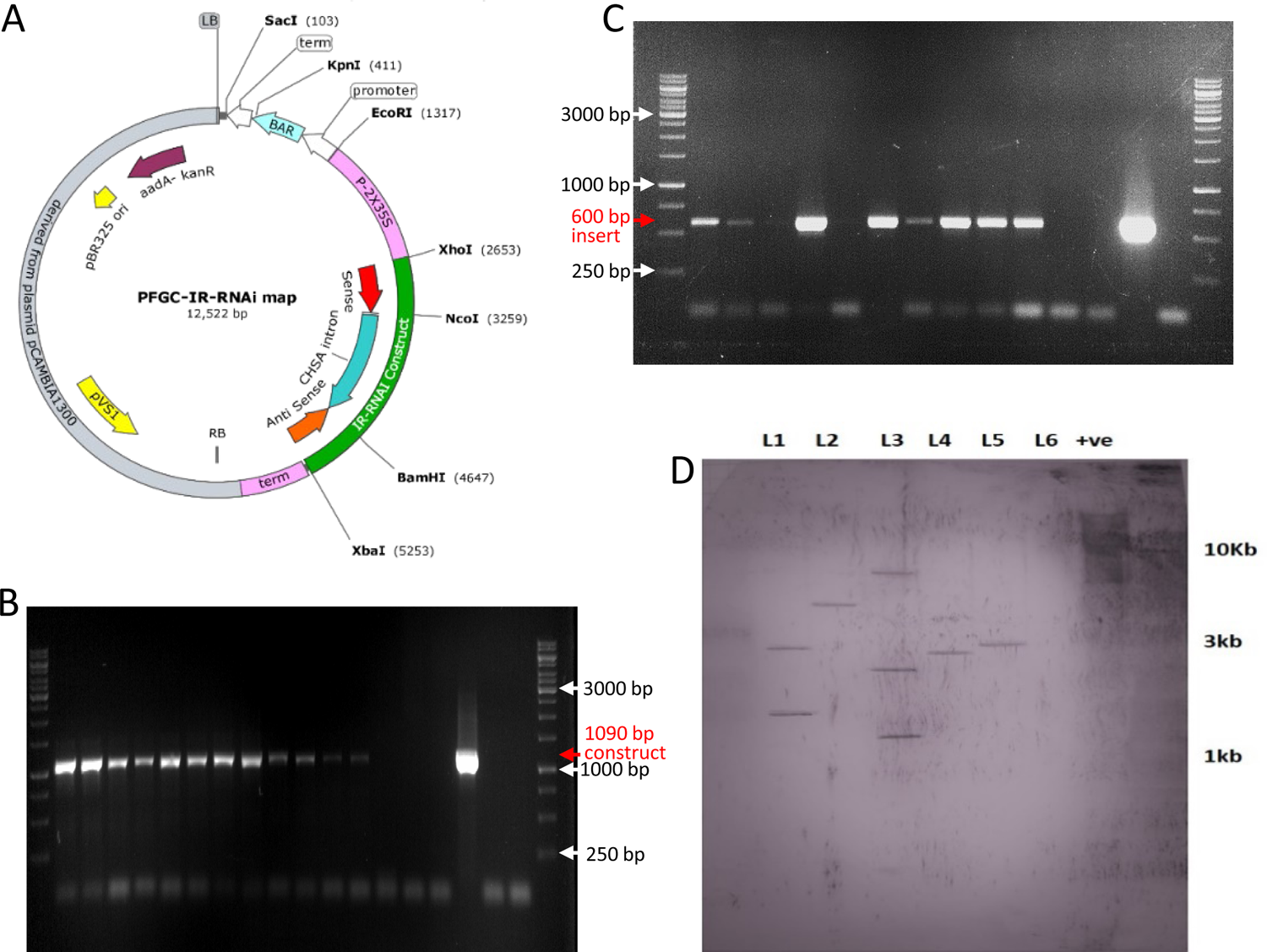
Development and confirmation of transgenic tobacco. (A) Transformed vector pFGCIR_RNAi, with schematic representation of the RNAi cassette containing sense-antisense orientation with fused gene fragments of *Myzus persicae* cathepsin L, *Bemisia tabaci* ecdysone receptor, and *Phenacoccus solenopsis* chitin synthase (B) Amplification from tobacco with construct-specific primers (1090 bp); Lane 1-12 (transgenic lines), 13-15 (non-transgenic), 16 (+ve control plasmid), 17 &18 (-ve control, H_2_O only), L = 1kb DNA ladder (C) Amplification from tobacco with gene specific primers (600bp); Lane 1-10 (transgenic lines), 11 & 12 (nontransgenic), 13 (+ ve control plasmid), 14 (-ve control water) and L = 1kb DNA ladder (D) Southern blot analysis of five transgenic tobacco plants confirmed by PCR. Forty µg genomic DNA from each transgenic plant were digested with EcoRI and probed with the PCR-amplified sense sequence. Lanes 1-5 transgenic lines; Lane 6 non-transgenic; Lane 7 +ve plasmid.

### 2.2. Agrobacterium-*mediated tobacco transformation*

The RNAi plasmid pFGC5941-IR_RNAi was transformed into LBA1404 (*Agrobacterium tumefaciens*) by electroporation. A single bacterial colony was confirmed with PCR and grown for 48 hours in LB medium with kanamycin (50 µg/ml) and rifampicin (25 µg/ml) at 160 RPM, 28^ꅶ^C. *Nicotiana tabacum* plants were transformed using the leaf disc method (Horsch et al., 1985). Murashige and Skoog medium (Murashige and Skoog, 1962) with kanamycin (50 mg/L), cefotaxime (250 mg/L), 1-naphthalene acetic acid (250 µg/L) and 6-benzylaminopurine (1 mg/L) was used for the selection of transformants. The transgenic plants were transferred to peat mix.

### 2.3. Transgene analysis

Genomic DNA was isolated from leaf tissue of transgenic and non-transgenic tobacco plants by cetyltrimethylammonium bromide (CTAB) method (Stewart and Via, 1993). All transgenic lines were confirmed by PCR-amplification with gene-specific and construct specific primers (Table S2). Southern blotting was performed to confirm the integration of target genes in *N. tabacum* plants. Forty µg of genomic DNA was digested with EcoRI restriction enzyme. Digested DNA was separated by electrophoresis on an 0.8% agarose gel and transferred onto nylon Hybond–N + membrane (Roche). The 600 bp sense sequence was used to make a digoxigenin-labeled probe. Hybridization and all other procedures were followed according to manufacturer’s instructions using the DIG High Prime DNA Labeling and Detection Starter Kit1 (Sigma Aldrich).

### 2.4. Aphid bioassays

All aphids were from a strain of red, tobacco-tolerant *M. persicae* (Ramsey et al., 2014) that was originally collected from tobacco by Stewart Gray (Cornell University). The aphid colony was maintained on tobacco plants in a growth chamber. T1 transgenic tobacco plants (lines 1, 2, 3, 4, and 5) and non-transgenic (control) were grown at 24±2°C with 70% humidity under a 17 hours light and 7 h dark cycle. All plants were used for insect bioassay at the 5-6 leaf stage. Five adult *M. persicae* per replication were released on the leaf surface. On the next day, all adult aphids were removed and five-first instar nymphs were remained on a leaf. Aphid mortality and fecundity was observed on daily basis. There were six replicates for each treatment.

### 2.5. Real-time PCR analysis

After 3, 5, and 8 days of feeding on transgenic plants, five aphids from each line were collected RNA isolation. RNA was extracted by using the RNA isolation Kit (SV Total RNA isolation system, Promega) and quantified by using NanoDrop 2000 Spectrophotometer (Thermo Fisher). Two hundred ng total RNA was used for first-strand cDNA synthesis using “Primescript First Strand cDNA Synthesis” kit, Takara, China. Power SYBR Green PCR Master Mix (Thermo Fisher) was used in 10 µl of reaction volume and analyzed with a Quantstudio 6 Real-time PCR system (Thermo Fisher Scientific) under the following PCR profile: 95°C for 3 minutes followed by 40 cycles of 95°C for 30 s, 56°C for 30 s and 72°C for 50 s. All reactions were performed in triplicate using gene-specific primers and actin primers as an internal control (Table S2).

### 2.6. Biochemical analysis of aphids

Fifteen mg of *M. persicae* were collected from transgenic line 3 and non-transgenic tobacco plants. Aphids were placed in a 2 ml microcentrifuge tube, with 180 µl of lysis buffer solution (100 mm KH2PO4, 1 mm dithiothreitol (DTT) and 1 mm ethylenediaminetetraacetic acid (EDTA), pH 7.4) and 5 2 mm stainless steel beads. Samples were crushed on a bead beater (1600 MiniG, Automated mini tissue homogenizer and cell lyser) for 30 sec.

A standard curve was calculated by using bovine serum albumin (BSA) according to manufacturer’s instruction (B6916, Sigma Aldrich). Total protein was analyzed from the aphid bodies using the Bradford method (Hammond and Kruger, 1988; Walker, 2002). Each homogenate sample was centrifuged at 180 *g* at 4°C for 5 min. Thirty-three µl of the supernatant was added to 967 µl of Bradford reagent (B6916: Sigma Aldrich) in a 1.5 ml microcentrifuge tube. The same concentration of BSA was also dissolved in Bradford buffer reagent and used as a standard. All samples were incubated at room temperature for 15-20 min. After incubation, protein concentrations were determined with a spectrophotometer at 595 nm (Foray et al., 2012).

According to an established protocol, D-glucose was used to make a standard curve (8418: ScienCell). Carbohydrates were analyzed using anthrone reagent and a colorimetric method (Van Handel, 1965). Aphid samples, homogenized as described above, were used for carbohydrate and glycogen extraction according to a previously published protocol (Foray et al., 2012; Van Handel, 1965). For carbohydrate analysis, 150 µl of the supernatant was taken from the homogenate after vortexing and centrifugation at 180 *g* at 4°C for 15 min. The supernatant was transferred to separate 1.5 ml microcentrifuge tube and evaporated at room temperature for 45 min, or until the volume had reached 40 µl. Then 960 µl of anthrone reagent (SKU319899: Sigma Aldrich) was added and samples were incubated at room temperature for 15-20 min. Samples were heated in water bath at 90°C for 15 min. Sample absorbance was measured at 625 nm in a spectrophotometer (BioMate 3S: Thermo Scientific) using D-glucose (G8270: Sigma Aldrich) as a standard.

For the glycogen assay a pellet was obtained from aphid homogenate after centrifugation as described above. The pellet was twice washed with 400 µl of 80% methanol, followed by vortexing and centrifugation at 16000 *g* for 5 min at 4°C. The supernatant was removed and 1 ml of anthrone reagent was added to the pellet. The mixture was then incubated for 15 min at 90°C in a water bath, followed by snap cooling in ice. The cooled mixture was filtered by using a 0.45 µm diameter low-protein filter membrane (SLHVM33RS: Fisher Scientific). The glycogen contents were analyzed at 625 nm in a spectrophotometer with D-glucose as the standard (Foray et al., 2012).

Total lipids were determined by using the vanillin assay procedure (Van Handel, 1985). Triolein (Y0001113: Sigma Aldrich) was used to make a standard curve. Vanillin reagent was prepared according to a standard protocol (Foray et al., 2012). One hundred µl of supernatant (after centrifugation of the aphid homogenate) was shifted to another microcentrifuge tube and heated at 90°C until complete evaporation. Ten µl of 98% sulphuric acid (H_2_SO_4_) was added to the tube and incubated at 90^ꅶ^C for 2 min in a water bath, followed by snap cooling in ice. One ml of vanillin reagent was added to the cooled mixture and incubated for 15 min at room temperature. Samples were shifted to cuvettes and analyzed for lipid detection at 525 nm absorbance in a spectrophotometer, with triolein as standard.

#### Protein and carbohydrate quantification in aphid honeydew

Transgenic lines 1 and 3 and non-transgenic control tobacco plants were used to detect protein and carbohydrate in the honeydew of aphids feeding on them. Ten first instar age-synchronized aphid nymphs were released on the leaf surface, with 15 replications of each treatment. To measure honeydew production, Whatman paper was placed in the cage horizontally in such a way that honeydew dropped onto the paper (Fig. S1). Aphids were allowed to feed for 24 h on plants, after which the filter paper was removed and soaked in 0.1% (w/v) ninhydrin solution. After incubation at 65°C for 30 mins, purple ninhydrin-stained spots corresponding to aphid honeydew appeared (Kim and Jander, 2007). For honeydew quantification, filter papers were cut into small pieces and placed in microcentrifuge tubes. One ml of 90% methanol was added to each tube and placed at 4^°^C for 1 h with continuous agitation. The solution was centrifuged for 1 min at 6,000 *g* and the supernatant was used to quantify honeydew by taking absorbance at 500 nm with 90% methanol as the standard (Nisbet et al., 1994; Rashid et al., 2017). Absorbance per honeydew spot was calculated.

For carbohydrate and protein analysis in honeydew of aphids, honeydew was collected as described above, but using aluminum foil instead of filter paper. The aluminum foil was washed with 500 µl of distilled water until all honeydew was collected. Individual samples were collected in separate microcentrifuge tubes. Forty µl of solution from each replication were separated and used for carbohydrate quantification by using anthrone reagent and absorbance at 625 nm, as described above. The remaining honeydew solution (460 µl) was used for the detection of protein with the ninhydrin method. The remaining honeydew solution was lyophilized and resuspended in 10 µl of sterile water. Each sample was spotted on Whatman filter paper and treated with 0.1% ninhydrin solution followed by incubation at 65^ꅶ^C for 30 min. Protein abundance was determined with a spectrophotometer at 500 nm as described above. There were twenty replicates for each treatment.

2.6 Coccinella septempunctata *bioassay*

First instar *C. septempunctata* larvae from a colony maintained by Todd Ugine (Department of Entomology, Cornell University) were used for bioassays. First-instar beetle larvae were fed ad libitum with ten aphids per day. After the second instar, each larva was given 30 mg aphids per day. There were three treatments with ten replications each. Two treatments involved aphids raised on transgenic tobacco (pFGC5941-IR_RNAi lines 1 and 3) and the third involved aphids that had fed on control plants. Larval growth was monitored on a daily basis. The collected data were larval mortality, larval weight, larval period, pupal period, pupal mortality, adult weight on emergence, and adult emergence rate.

### 2.7. Statistical analysis

Statistical comparisons were done using JMP 13 (https://www.jmp.com).

## 3. Results

### 3.1. Development of transgenic tobacco lines

Based on multiple alignments of target genes, 200 bps of the highly conserved region of *M. persicae* cathepsin L (AJ496197), *B. tabaci* ecdysone receptor (EF174329), and *P. solenopsis* chitin synthase (KF384512) were selected and cloned in the pFGC5941 RNAi vector. The resulting plasmid was named as pFGC5941-IR_RNAi (Fig. 1A; Table S1). Based on BLAST comparisons to the *M. persicae* genome (Mathers et al., 2017), only the cathepsin L part of the construct will inhibit gene expression in this species. The other two genes, ecdysone receptor and chitin synthase, do not have lengths of 20 nucleotides or more identity to the aphid genome and thus are not expected to initiate cleavage reactions. Comparisons to available sequence data of *C. septempunctata* and other coccinellid beetles in GenBank showed no extensive sequence identity, *i.e.* no stretches of identity greater than 20 nucleotides, which would be required for expression silencing by RNA interference in the beetles.

Tobacco lines were transformed with pFGC5941-IR_RNAi using *Agrobacterium tumefaciens* and confirmed by PCR with gene-specific and construct-specific primers (Fig.1B, C). Five independent transgenic lines were selected for further assays. A Southern blot was performed by using digoxigenin-labeled sense sequence as a probe to confirm the integration of target genes. The number of integration sites ranged from one to three and integration of all three target genes in the tobacco genome was confirmed (Fig. 1E).

### 3.2. Aphid responses to cathepsin L RNAi

Real-time quantitative PCR was used to analyze the cathepsin L mRNA levels after 3, 5, and 8 days of aphid feeding, showing significant knockdown of cathepsin L expression in aphids feeding on all transgenic tobacco lines as compared to control (Fig. 2). With the exception of line 5, silencing efficiency increased over time from day 3 to day 8. The cathepsin L expression level in aphids after 8 days on transgenic lines 3 and 4, was less than 5% of that in aphids the control plants.

**Figure 2.**
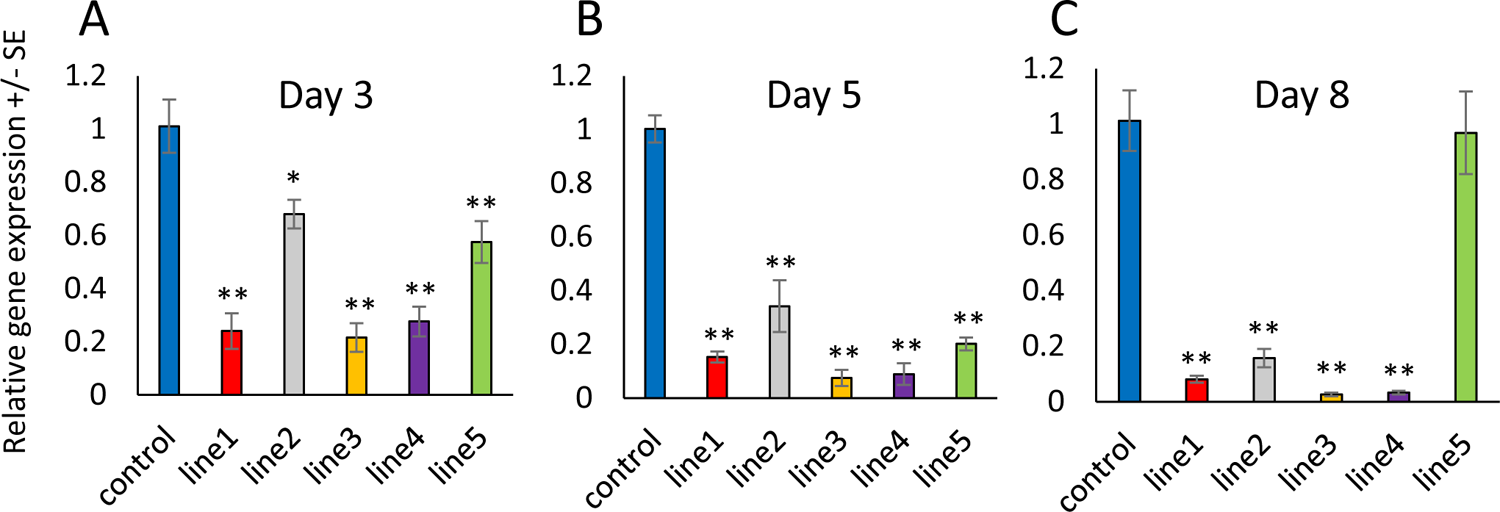
Expression profiling based on qRT-PCR of cathepsin L in aphids feeding on tobacco plants with and without pFGC-IR_RNAi constructs. (A) Relative expression after three days (B) Relative expression after five days days (C) Relative expression after eight days. Mean ± SE of N = 3, *P < 0.05, **P < 0.005, Dunnett’s test relative to control at the same time point.

Given the significant reduction in cathepsin L gene expression, a bioassay was conducted to determine whether this affects the survival and fecundity of *M. persicae*. All five transgenic lines showed significantly increased aphid mortality on day 12 compared to the control (P < 0.05, Dunnett’s test; Fig. 3A), with the highest mortality (68% and 80% respectively) observed in transgenic lines 1 and 3. Aphid fecundity was significantly reduced on all days in aphids fed on transgenic line 3 compared to the control (P < 0.05, *t*-test; Fig. 3B). Additionally, in line 3 the first nymph production was observed on day 8, whereas in all other lines nymph production started two days earlier (Fig. 3B). Total nymph production by day 12 was significantly decreased in all transgenic lines as compared to the control (P < 0.05, Dunnett’s test; Fig. 3C).

**Figure 3.**
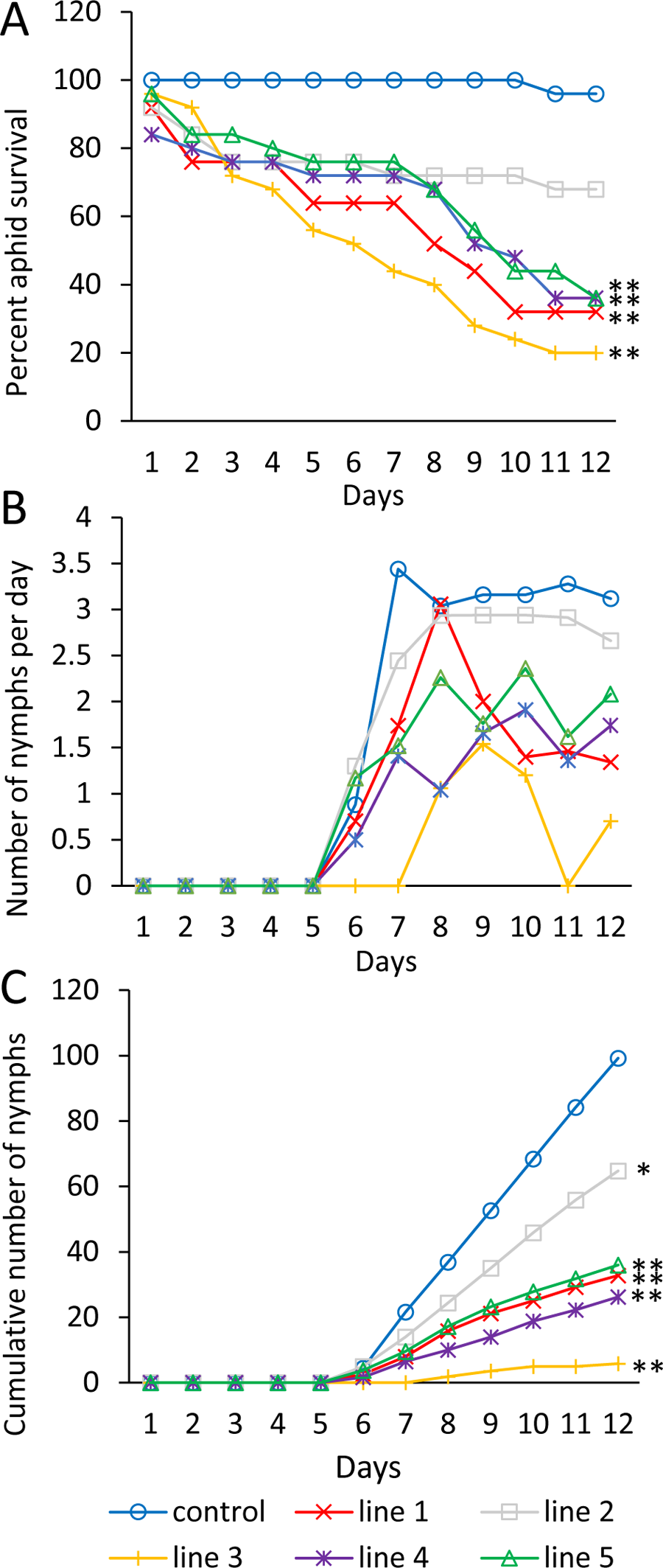
Aphid growth on transgenic and control tobacco lines (A) Percent survival of adult aphids feeding on transgenic and non-transgenic plants, (B) Nymphs produced per adult aphid per day (c) total number of nymphs produced on each tobacco line. Mean of N = 5, *P < 0.05, **P < 0.005, Dunnett’s test relative to control on day 12.

Based on the known function of cathepsin L as a gut protease, we hypothesized that expression silencing could reduce nutrient uptake by the aphids. Transgenic line 3, which showed lowest level of cathepsin L expression (Fig. 2) and the greatest reduction in survival and reproduction of aphids (Fig. 3) was analyzed to determine total protein, carbohydrates (sugar and glycogen), and lipids in whole aphid bodies. This showed significantly reduced levels of protein and glycogen in aphids fed on transgenic line 3 as compared to aphids fed on non-transgenic plants (Fig. 4A, B). Conversely was a significant increase in sugar and total lipids in aphids on transgenic plants relative to controls (Fig. 4C, D).

**Figure 4.**
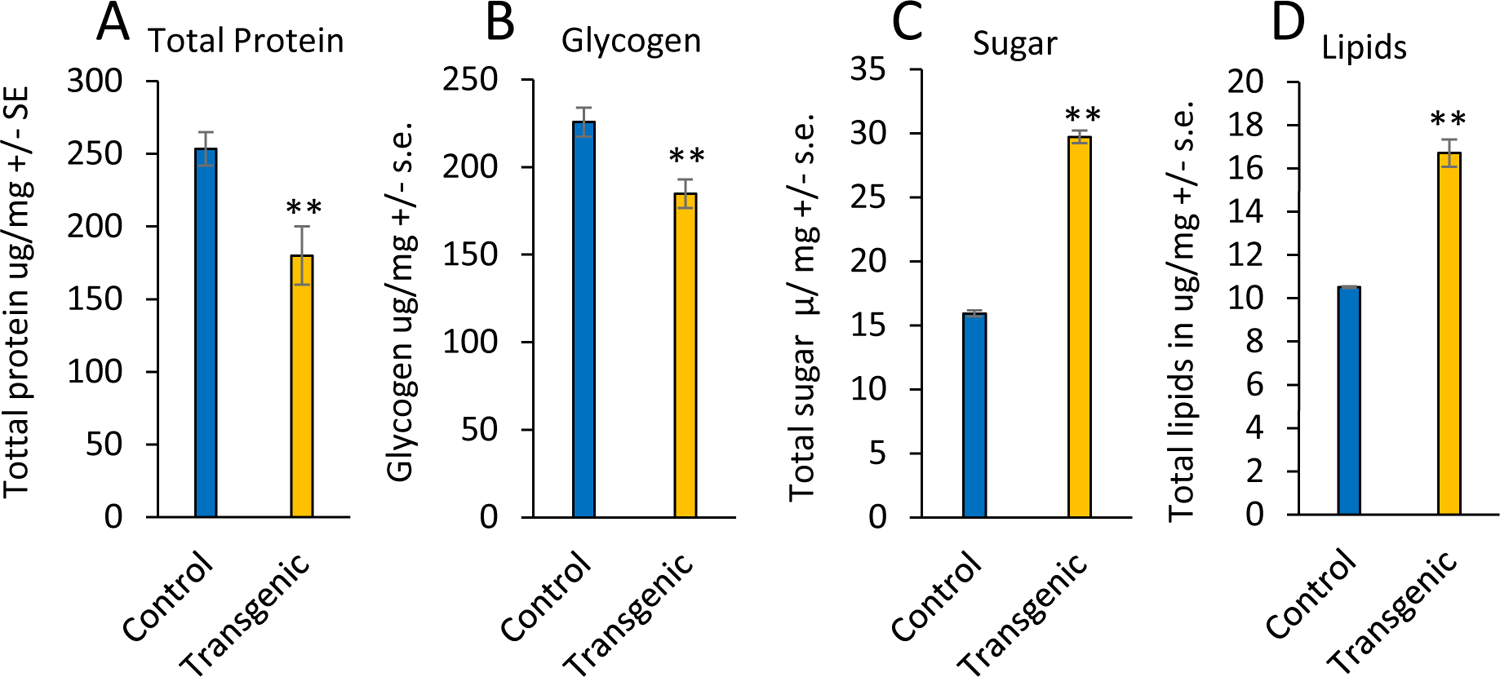
Biochemical analysis of aphids fed on transgenic and non-transgenic tobacco plants. (A) Total protein content of aphids, (B) Glycogen content of aphids, (C) Total sugar content of aphids, (D) Total lipid content of aphids. Mean ± SE of N = 3. **P < 0.05, two-tailed *t*-test.

The lower protein content in the bodies of aphids on transgenic plants (Fig. 4A), suggested less protein uptake from the phloem sap during passage through the gut and therefore increased protein excreted in the honeydew. Assays of aphid honeydew production by measuring ninhydrin-stained spots on filter paper (Fig. S1) showed that aphids produced fewer honeydew spots (Fig. 5A), but with more protein per spot (Fig. 5B) on the transgenic lines 1 and 3 compared to controls. Separate measurement of honeydew deposited on aluminum foil showed a significantly larger amount of protein in the honeydew of aphids feeding on transgenic lines 1 and 3 compared to those feeding on control plants (Fig. 5C). In contrast, the carbohydrate level was significant lower in aphids feeding on the transgenic lines (Fig. 5D). Together, these measurements show reduced honeydew production and a higher protein/carbohydrate ratio in the honeydew from aphids feeding on transgenic lines 1 and 3, consistent with the hypothesis that the aphids have less efficient amino acid uptake after cathepsin L expression silencing.

**Figure 5.**
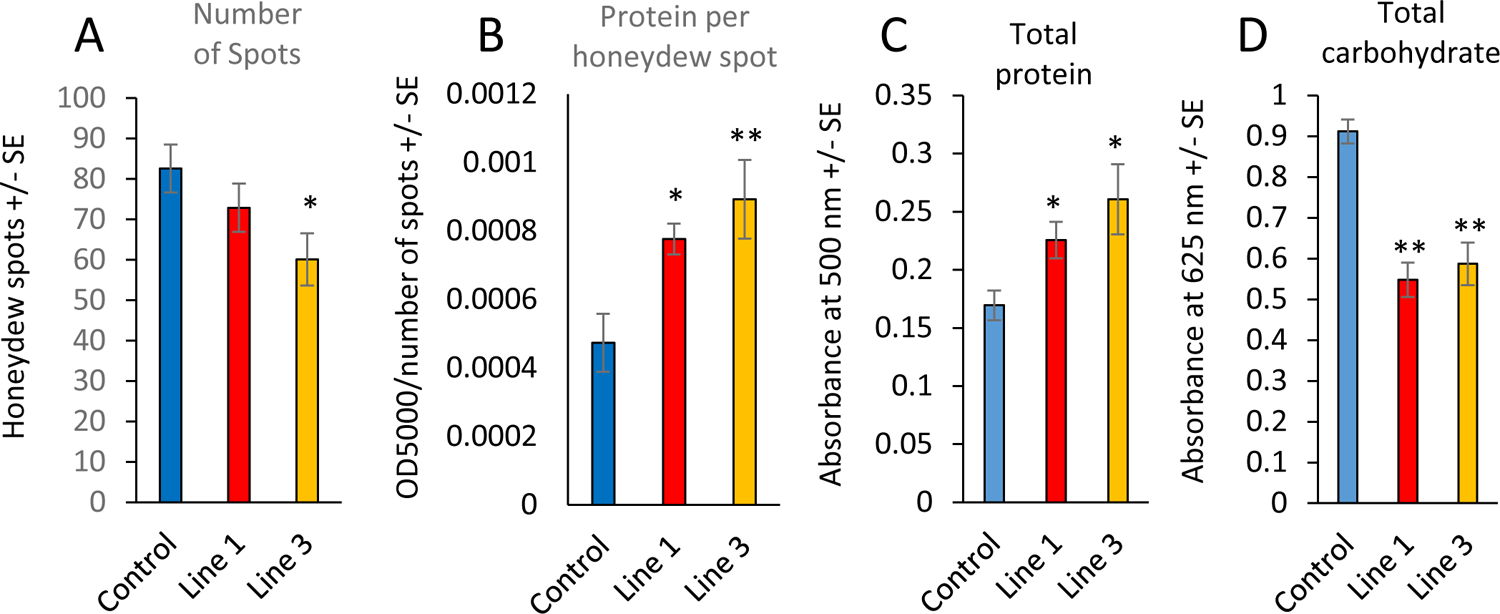
Quantification of protein and carbohydrate ratio in honeydew of aphids feed on transgenic and non-transgenic tobacco plants. (A) Number of honeydew spots produced by 10 aphids in 24 hours, mean +/− SE of n =17, (B) Protein content per honeydew spot, as measured by ninhydrin staining, mean +/− SE of n = 17, (C) Total honeydew protein, as measured by ninhydrin staining and absorbance at 500 nm of honeydew collected on aluminum foil, mean +/− SE of n =20, (D) Total honeydew carbohydrate, as measured by anthrone reagent staining and absorbance at 625 nm of honeydew collected on aluminum foil, mean +/− SE of n =20. *P < 0.05, **P < 0.05, Dunnett’s test relative to control.

### 3.3. *Growth of* Coccinella septempunctata

Given the decreased protein content of aphids feeding from transgenic lines 1 and 3, we hypothesized that they would be less suitable food material for predatory insects. Bioassays conducted with *C. septempunctata* showed that beetle larvae consuming aphids that had fed on the transgenic tobacco had a significantly lower body mass after nine days compared to those consuming aphids from control plants (Fig. 6A). There was a significant lengthening in the *C. septempunctata* the pupal period, but not the larval period, when feeding on aphids from transgenic plants (Fig. 6B, C). Beetles consuming aphids from transgenic plants had a significantly reduced pupal mass compared to controls (Fig. 6D). However, there was no significant difference in the rate of adult beetle emergence (Fig. 6E).

**Figure 6.**
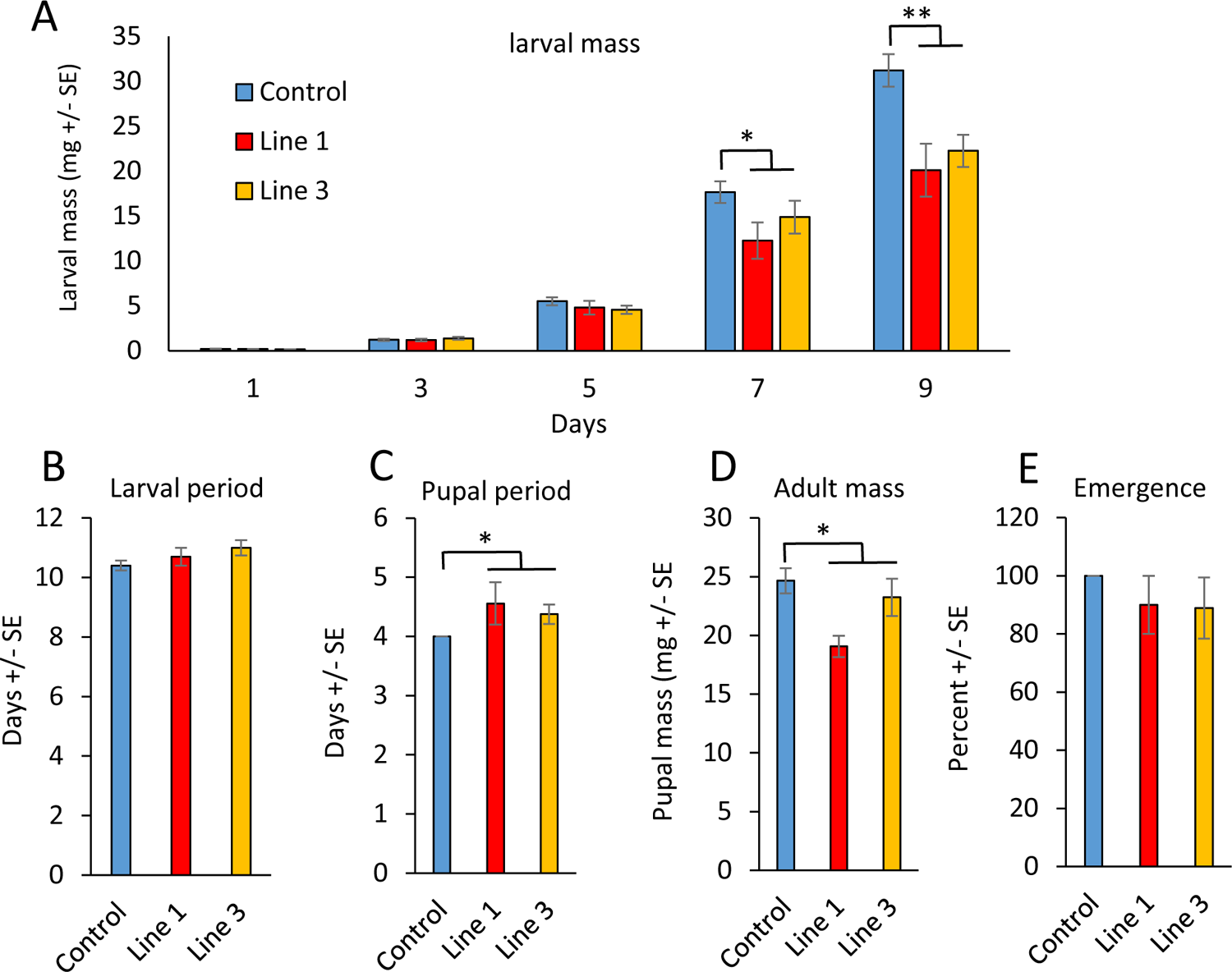
Performance of Coccinella septempunctata, feeding on Myzus persicae that were raised on RNAi transgenic tobacco lines 1 and 3, and non-transgenic control plants. (A) larval mass over nine days, (B) time spent as larvae, (C) time spent as pupae, (D) pupal mass, (E) percent emergence. Mean ± SE of N = 10, *P < 0.05, **P < 0.005 Wilcoxon rank sum test.

## Discussion

The introduction of novel and durable control strategies is critical for sustainable insect pest management. RNAi-based plant production opens a new era in the eco-friendly management of sucking insects (Bhatia and Bhattacharya, 2018; Luo et al., 2017; Poreddy et al., 2017; Raza et al., 2016). Silencing of target genes in insects mainly depends upon the route of delivery of dsRNA and the function of the targeted gene (Huvenne and Smagghe, 2010). Therefore, cathepsin L, a cysteine protease that functions in aphid guts is an attractive target for plant-mediated RNAi. Our observation that cathepsin L expression knockdown (Fig. 2) can cause up to 80% mortality, reduced fecundity, and a delay in production of nymphs (Fig. 3) confirms the choice of this gut-expressed gene as an efficacious target for RNAi knockdown.

The decreased amino acid content in aphids feeding on cathepsin L RNAi plants is consistent with the known role for this enzyme in digesting protein in the aphid gut, thereby making free amino acids available for uptake into the insect body. Although the fole of cathepsin L as an aphid gut protease is well-established, injection of dsRNA into *A. pisum* bodies suggested additional functions in aphid physiology (Sapountzis et al., 2014a). In *Helicoverpa armigera* (cotton bollworm), cathepsin L was shown to be required during moulting (Liu et al., 2006). Thus, we cannot completely rule out indirect effects of cathepsin L silencing in other parts of the aphid body on protein degradation and amino acid uptake from the gut. However, prior research has shown that silencing of aphid gut enzymes by plant-mediated RNAi is efficient (Tzin et al., 2015) and, in particular, ingested cathepsin L dsRNA silences expression of this gene in the aphid gut rather than in other body parts (Sapountzis et al., 2014a).

In some insect species, compensatory feeding behavior has been observed on diet with reduced nutrient content. For instance, nitrogen uptake by *Spodoptera exigua* caterpillar feeding on *Nicotiana attenuata* is subject to inhibition by protease inhibitors from the host plants (Steppuhn and Baldwin, 2007). The caterpillars compensate for the protease inhibitors by consuming more material from wild type *N. attenuata* than from mutant plants that lack this enzyme. However, aphids feeding on plants expressing cathepsin L RNAi constructs produced fewer honeydew spots (Fig. 5A) and had less total sugar output in 24 hours (Fig. 5D) than those on control plants, suggesting that they are consuming less phloem sap and are therefore not compensating for reduced protein uptake. Although prior experiments have demonstrated variation in aphid honeydew composition (Faria et al., 2008; Fischer et al., 2002), to the best of our knowledge the current study is the first to show specific effects of an RNAi construct.

Cathepsin B, an expanded family of gut-expressed aphid proteases (Pyati et al., 2011; Ramsey et al., 2007; Rispe et al., 2008), might partially compensate for the absence of cathepsin L. Similar to the negative effects of silencing cathepsin L (Fig. 3), silencing several cathepsin B genes resulted in reduced aphid growth (Mathers et al., 2017). Simultaneous knockdown of cathepsin L and one or more cathepsin B genes might result in synergistic effects on protein digestion and even greater negative effects on aphid performance than either construct alone. Similar synergistic effects were observed when silencing expression of multiple osmoregulatory gene in *M. persicae* guts (Tzin et al., 2015).

Due to the high specificity of dsRNA, there is a reduced chance of adverse effects on non-target arthropods. However, even in the absence of gene expression knockdown in predatory insects by RNAi, there can be indirect effects in the predator-prey interaction. In particular, a change in the nutritive value of aphids can influence interactions with predators and parasitoids. For instance, variation in dietary carbohydrate can produce quantitative effects on the growth of a larval parasitoid (Thompson, 1982). Consistent with a prior observation that *C. septempunctata* attained more weight when fed on a high-protein (Atwal, 1963), beetle larvae consuming low-protein aphids from cathepsin L RNAi plants grew more slowly (Fig. 6A) and produced smaller adults (Fig. 6D). This reduction in *C. septempunctata* growth could have negative effect on the populations of this beneficial insect. However, in natural settings, *C. septempunctata* are undoubtedly confronted with prey species that varying nutritional value. It is possible that, similar to caterpillars on *N. attenuata* (Steppuhn and Baldwin, 2007), C *. septempunctata* larvae may compensate for a low-protein diet by consuming more aphids. If this is the case, integrated pest management in an agricultural setting may actually be improved if aphids have a lower protein content.

Together, our results confirm the effectiveness of gut-localized aphid cathepsin L as a target for plant-mediated RNAi. It is likely that similar RNAi constructs can be used to ameliorate *M. persicae* infestation of cotton and other important crop plants. However, the observation of reduced *C. septempunctata* growth when consuming aphids from cathepsin L RNAi plants suggests that, during the regulatory approval process of such transgenic crops, it will be necessary to consider more than just direct negative effects on gene expression in beneficial insects.

## Acknowledgments

IR was funded by the Higher Education Commission, Government of Pakistan to carry out research at the National Institute of Biotechnology and Genetic Engineering and the Boyce Thompson Institute under IRSIP (International Research Support Initiative Program). Research relating to RNAi effects on predators was funded by United States Department of Agriculture award G14825 R44 01337 to GJ. We thank Todd Ugine for providing *Coccinenella septempunctata.*

## Supplementary materials

**Supplemental Table S1.** Inserted DNA sequences

**Supplemental Table S2.** Primers used in this study

**Supplemental Figure S1.** Experimental setup

## Supplementary Data

### Supplemental Table S1.

Selected sequences (200 bp each) from ecdysone receptor (EF174329), cathepsin L (AJ496197) and chitin synthase (KF384512).

**Gene 1 = Ecdysone Receptor (*Bemisia tabaci*)**

~~~
aaggatagttaatgctgcaccagaagaagaaaatgtagctgaagaaaggtttaggcatattacagaaattacaattctcactgtacagttaattgtggaattttctaagcgattacctggttttgacaaactaattcgtgaagatcaaatagctttattaaaggcatgtagtagtgaagtaatgatgtttagaatggcaa
~~~

**Gene 2 = Cathepsin L (*Myzus persicae*)**

~~~
tggtgtgctagtatctttgagtgagcaaaaccttattgattgctctcgtaaatacggtaacaatggttgtgaaggtggtcttatggatcttgcattcaagtacatcaaatctaacaagggactagatactgaaaaatcttacccatatgaagccgaggatgacaagtgccgttataatcctgataactctggtgccactg
~~~

**Gene 3 = Chitin synthase (*Phenacoccus solenopsis*)**

~~~
tacgctattggtcactggctgcaaaaagctaccgaacatatgatcggatgtgtactttgtagtcccggatgcttttcgttgtttcgtggtaaagctctgatggatgataatgttatgcgcaaatataccactaaatctcaagaagctagacattatgtgcaatacgatcaaggagaggatcgttggctgtgtacattact
~~~

### Supplemental Table S2.

List of primers used for transgenic confirmation.

**Gene Specific Primers**

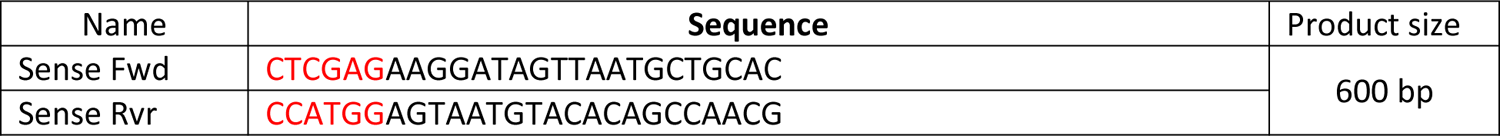

**Construct Specific Primers**

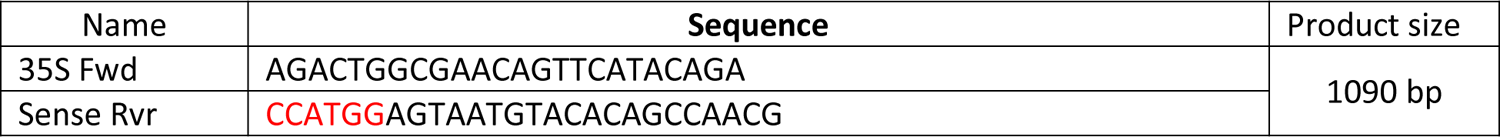

**Supplemental Figure S1.**
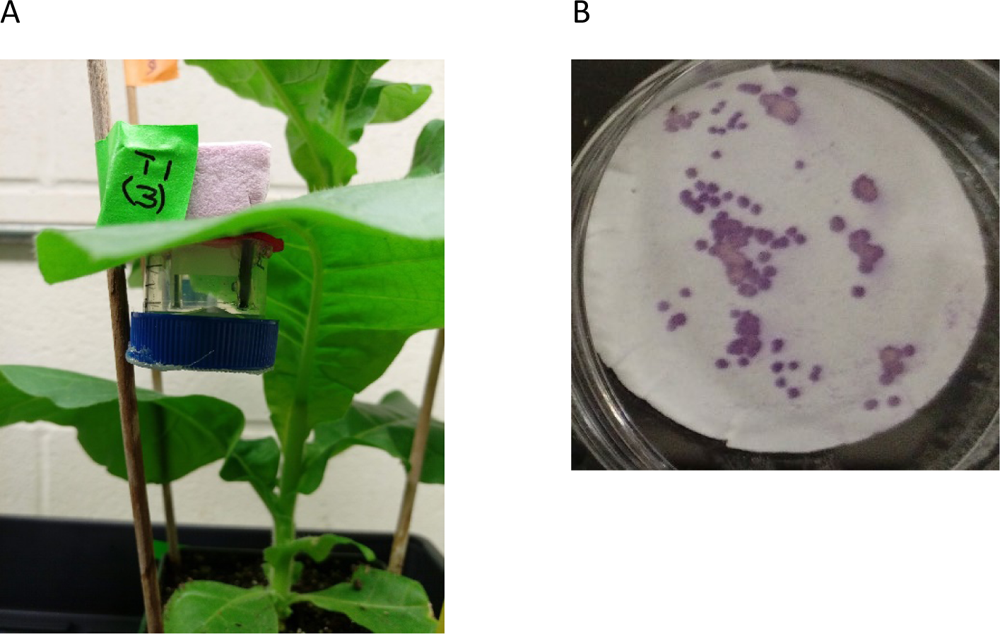
(A) Collection of aphid honeydew on Whatman filter paper that was placed at the bottom of insect cages, and (B) representative staining of honeydew spots on filter paper with ninhydrin.

